# Stress-Induced Mechanical Memory in Respiratory Mucus: Anisotropy, Network Reorganization, and Directional Transport

**DOI:** 10.64898/2026.07.16.738946

**Authors:** Ameya G. Prabhune, Behnam Rezaei, Andy S. García-Gordillo, Moumita Das, Franck J. Vernerey, Nuris Figueroa-Morales

## Abstract

Mucus transport is essential for lung health, as ciliated cells constantly propel mucus outward to clear bacteria, viruses, and particles. This defense relies on a material that must be elastic enough to maintain ciliary traction, but capable of reorganizing under sustained directional loading. How the mucin network reconciles these demands, and whether it retains a memory of the stresses it experiences, remains poorly understood. Here we show that lung mucus develops a persistent, direction-dependent mechanical asymmetry under physiologically relevant stress—a mechanical memory encoded in the slow scaffold of the network. Using bulk rheology, we find that directional pre-stress produces a residual anisotropy that grows with stress magnitude and persists long after the load is removed. A transient network model attributes this memory to a separation of timescales between transient bonds and a long-lived crosslink scaffold, and particle-tracking microrheology confirms that the memory reorganizes the network geometry at the scale of biological particles, biasing tracer diffusion along the axis of applied stress. The stresses required to induce memory are within the range generated by ciliary beating and remain below the mucus yield threshold, suggesting that mucociliary clearance operates in a regime where directional alignment accumulates without compromising the coherence of the mucus layer. This proximity to yield may not be coincidental, it allows mucus to accumulate mechanical memory under physiological forcing while remaining poised to flow during clearance events such as coughing.

---

The respiratory tract is lined by a thin mucus layer that continuously traps inhaled pathogens and pollutants, transporting them out of the lungs via the periodic coordinated beating of cilia [1–3]. To achieve efficient clearance, mucus must satisfy two seemingly contradictory mechanical demands: it must maintain solid-like coherence so microscopic ciliary tips can effectively grip and propel the bulk layer, while simultaneously retaining the capacity to flow and structurally adapt under sustained directional stress. Neither classical rheological limit suffices; a purely viscous fluid cannot form a cohesive moving layer, whereas a conventional elastic solid would resist continuous transport. Consequently, the mucus network relies on a dynamic viscoelastic architecture that balances short-timescale structural integrity with long-timescale directional adaptability. While this balance describes a stable material, it remains unknown whether repetitive ciliary forces drive a cumulative, non-equilibrium reorganization that alters how the network responds to subsequent driving.

In the lung, the mechanical loads driving mucus’ potential reorganization are inherently directional and repetitive. Ciliary beating, gravity, and coughing impose shear stresses with a preferred orientation, ranging from fractions of a Pascal during tidal breathing to tens of Pascals during ciliary metachronal activity, and hundreds of Pascals during coughing [4–6]. Prior work demonstrated that high mechanical stress can align the mucin network and rectify bacterial trajectories [7]. However, because the bidirectional feedback between physical forcing and network architecture remains unresolved, the temporal persistence of this alignment, its dependence on stress magnitude, its molecular origin, and its microscopic consequences for particle mobility remain unknown.

Mucus reconciles these dynamic demands through its composition as a complex viscoelastic material, consisting primarily of water and a supramolecular network of mucin glycoproteins cross-linked by a hierarchy of bonds with varying strengths and lifetimes. This network spans from strong, long-lived covalent disulfide bridges at the termini of mucin chains, to weaker transient associations, including hydrophobic interactions, calcium-mediated ionic bridges, hydrogen bonds, and physical entanglements [8–13]. This bond hierarchy gives rise to a broad spectrum of relaxation times: on short timescales, the network behaves as a soft elastic solid with a storage modulus nearly independent of frequency; on longer timescales, stress relaxes as transient bonds break and reform, eventually allowing macroscopic flow [14–16]. Crucially, the long-term structural consequences of this multi-scale relaxation spectrum under persistent, directional loading remain uncharacterized.

Here, we combine bulk rheology, particle-tracking microrheology, and transient network theory [17–19] to characterize the mechanical response of lung mucus to sustained and cyclic directional loading at physiologically relevant stress magnitudes. We show that directional stress produces a mechanical asymmetry that persists long after the load is removed—a structural memory encoded within the slow scaffold of the network. The magnitude of this memory scales with the applied prestress and is sufficient to bias microscopic particle diffusion at length scales relevant to inhaled pathogens and therapeutic vehicles. A constitutive rheological model, describing mucus as a transient network with stress-dependent bond attachment-detachment kinetics, successfully captures both the emergence and persistence of this anisotropy and maps it to the underlying crosslink dynamics. Together, these results reframe mucus not as a passive, isotropic barrier, but as a material endowed with mechanical memory in which the architecture retains and expresses a directional history of applied stress, with fundamental implications for mucociliary clearance, pathogen navigation, and targeted drug delivery.

## RESULTS AND DISCUSSION

### A. Directional stress produces a persistent mechanical asymmetry

We collected cow tracheal and lung mucus from excised lungs, stored it at 5 ^*°*^C, and performed all experiments within three days after butchering (see Methods). To characterize the mechanical response of mucus under directional loading, we applied a cyclic shear strain sequence oscillating between *γ* = −1 and *γ* = +1 (100%) [20], with each cycle following a sequence 0 → +1 → −1 → 0 over 40 s (Fig. 1A). Eleven cycles were applied to eliminate directional bias and allow the stress response to converge to a reproducible steady state (Fig. 1A–B). We take this stabilized 11th cycle as the baseline mechanical response of the network. The rapid convergence likely reflects the shear-driven reorganization of transient crosslinks and entanglements into a stable configuration. While absolute rheological values vary across samples with donor and environmental conditions, the qualitative stress–strain response is highly reproducible (Fig. S1), with a storage modulus exceeding the loss modulus, consistent with previous reports [7, 10, 21].

**FIG. 1.**
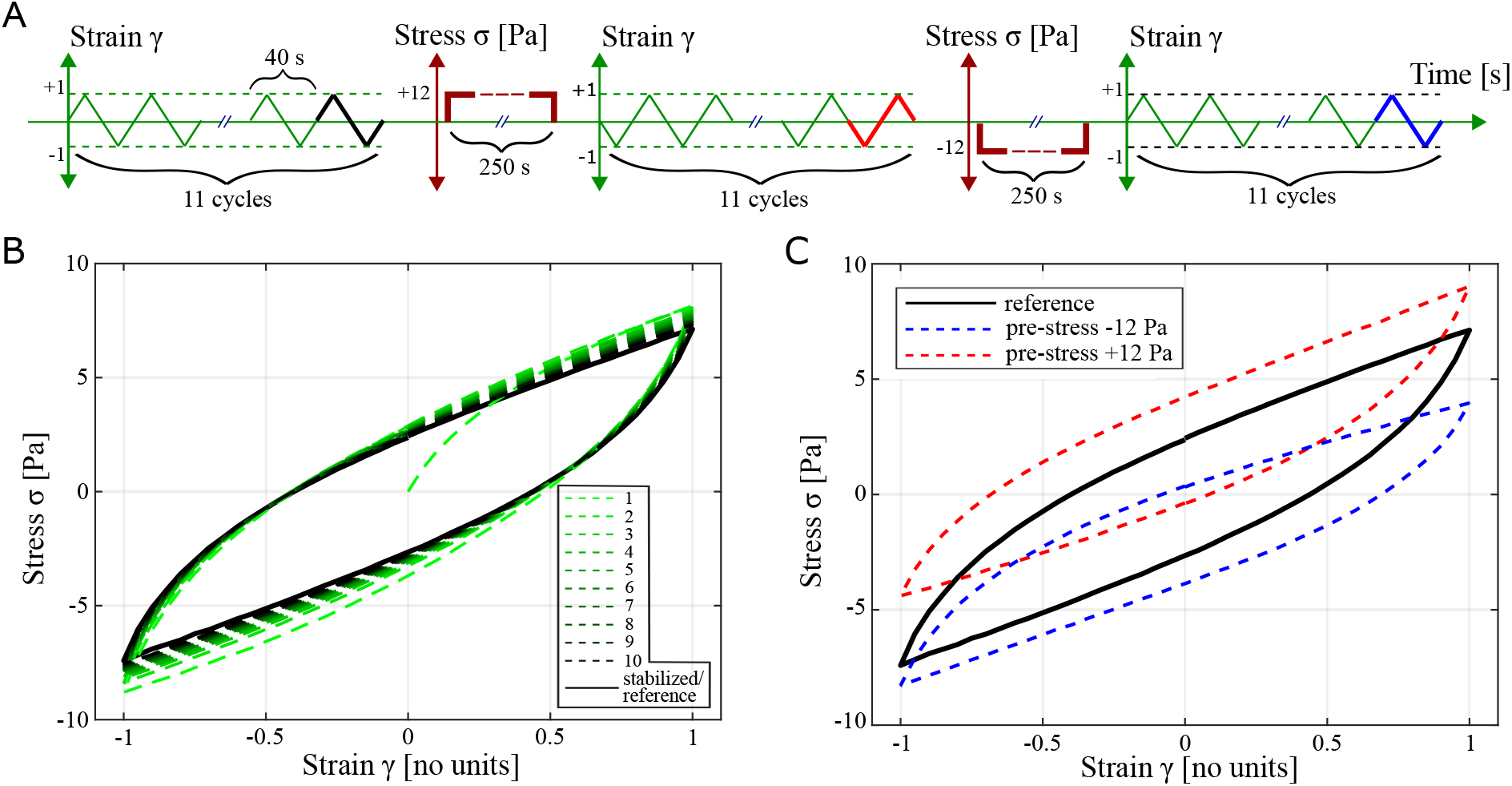
The effect of pre-stress on the mechanical behavior of cow tracheal mucus. (A) A series of 11 strain cycles were applied to the mucus sample, following the sequence 0 *→* +1 *→−* 1*→* 0, with each cycle lasting 40 s. The stress–strain response from the final cycle was used as the reference hysteresis behavior of the mucus. Subsequently, a pre-stress of 12 Pa was applied, in an arbitrarily determined positive direction, and the 11 strain cycles were repeated. The stress–strain response from the final cycle was analyzed to quantify the stressed behavior of the mucus. Finally, this procedure was repeated with pre-stress applied in the opposite (negative) direction. (B) Stress vs strain response of mucus for 11 cycles. The curve denoted in black is used as the reference response. (C) Comparison between the reference response of mucus and in the presence of *±*12 Pa pre-stresses.

To probe whether the network retains a directional memory of applied stress, we subjected the sample to a constant pre-stress of ± 12 Pa for 250 s before repeating the cyclic protocol. All experiments were conducted at strain rate 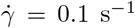, below the yield strain rate 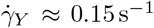 (Fig. S2, S3), ensuring that the network remained in the elastic regime and that pre-stressing did not cause irreversible flow. The key result is shown in Fig. 1C. The hysteresis curve shifts systematically upward or downward depending on the direction of the applied pre-stress. Moreover, the shift increases with the pre-stress magnitude (Fig. 2A). This direction-dependent residual offset is the signature of a mechanical memory: the network retains an elastic stress imbalance even after the pre-stress is removed, which we interpret as a slow-relaxing structural deformation of the polymer mesh aligned along the pre-stress direction.

**FIG. 2.**
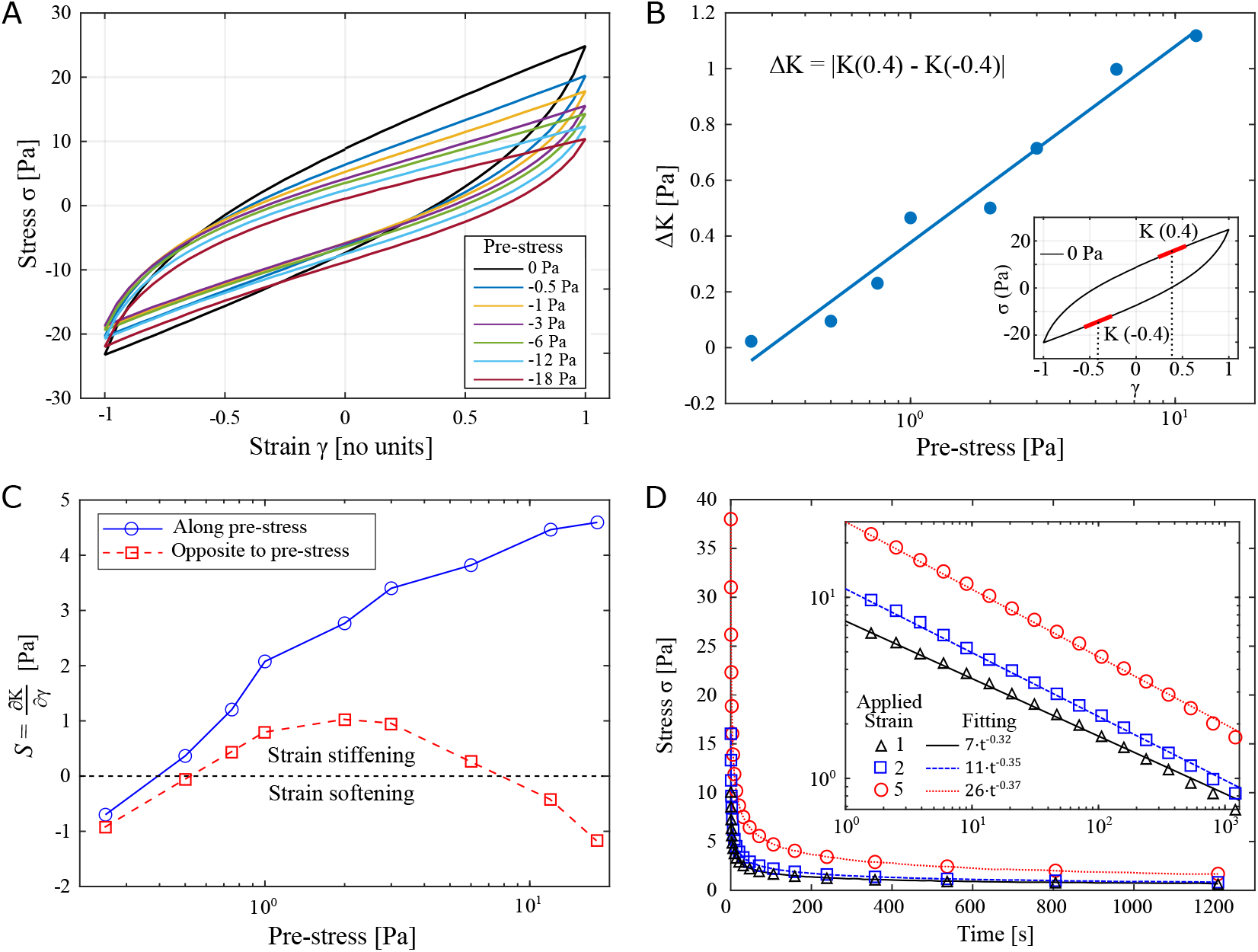
Macro-rheological signatures of shear-induced mechanical anisotropy and multi-scale stress relaxation. (A) Stress-strain Lissajous curves for several pre-stresses ranging from 0 Pa to -18 Pa. (B) Δ*K* quantifies the degree of mucus alignment, which increases with greater pre-stresses. (C) The strain stiffening parameter 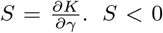 corresponds to strain-softening, and *S >* 0 to strain-stiffening. While the strain-stiffening effect is consistent for mucus along the direction of pre-stress, the opposite side shows a small strain-softening effect at higher pre-stresses. (D) Stress relaxation experiments for three applied strains (1, 2, and 5) and corresponding fittings show a broad range of relaxation times.

To quantify how this memory manifests itself as mechanical anisotropy, we computed the differential shear modulus *K*(*γ*) = *dσ/dγ* (the slope of stress-strain curves), separately for strains along the pre-stress direction (*γ <* 0) and against it (*γ >* 0). While *K* remains relatively stable along the pre-stress direction, it decreases monotonically under opposing loads (Fig. S5), evidencing a directional asymmetry in stiffness. We quantify this asymmetry by the modulus difference Δ*K* =| *K*(*γ* = +0.4) | −*K*(*γ* = −0.4), which grows monotonically with pre-stress magnitude (Fig. 2B), signaling a progressive transition from an isotropic gel to a directionally dependent medium.

This asymmetry is further captured by the strain-stiffening parameter *S* = *∂K/∂γ* (Fig. 2C). Along the pre-stress direction, *S* remains positive across all prestress magnitudes, indicating consistent strain stiffening. In the opposite direction, *S* transitions to negative values once the pre-stress exceeds approximately 8 Pa, indicating strain softening. We interpret this asymmetric response as a consequence of structural alignment: under directional loading, mucin fibers stretch along the stress axis and reach a high-tension state that resists further deformation. When strain is reversed, these aligned fibers buckle or shed transient crosslinks, producing the observed softening. Stress relaxation experiments for three applied maximum strains (1, 2, and 5) (Fig. 2D) reveal a broad spectrum of relaxation times.

Together, these results establish that physiologically relevant directional stresses imprint a persistent mechanical anisotropy on the mucin network. This directional memory has a magnitude and sign that depend on the history of applied load.

### B. Structural memory biases microscopic particle transport

While bulk rheology measures the average mechanical stiffness of the mucus layer, it cannot distinguish between a network that has simply hardened uniformly and one that has physically reorganized its microscopic structure. Particle-tracking microrheology allows us to resolve this distinction. If the mucin network genuinely aligns under stress, the spaces or “cages” within the mesh will stretch into elongated spaces, forcing embedded particles to diffuse primarily along the axis of alignment rather than in random directions (Fig. 3A). To test this, we used a custom rheo-microscopy setup that pairs a rheometer with an inverted microscope, allowing us to capture high-resolution video of fluorescent tracer particles while simultaneously applying controlled shear stress (Fig. S6). We tracked the microscopic trajectories of 200 nm beads in mucus that had been pre-stressed at 60 Pa for 250 s, and compared them to unstressed controls. The beads were coated with polyethylene glycol (PEG) to shield them from chemical binding or sticking to the mucin fibers, ensuring their movement purely probed the physical spacing of the network [14]. Furthermore, the diameter of the 200 nm beads is comparable to the size of respiratory viruses [22] and potentially therapeutic particles [23], making this length scale highly relevant for human health.

**FIG. 3.**
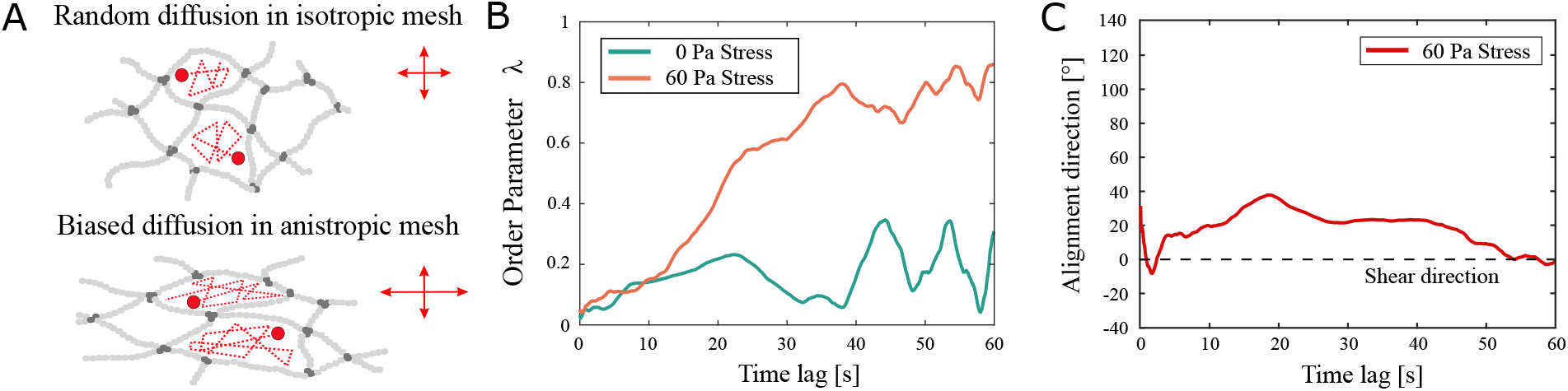
Biased particle diffusion in anisotropic mucus. (A) Sketch representing the trajectories of tracers in isotropic vs anisotropic (elongated) cages. (B) Order parameter from the diffusion of microparticles in pre-stressed and non-stressed mucus. The pre-stressed mucus shows biased diffusion, consistent with an anisotropic mesh. (C) Maximum diffusion direction with respect to shear, obtained from the eigenvalue of the anisotropy tensor.

To quantify this directional bias in particle motion, we analyzed the displacement covariance matrix, *C*_*ij*_(*t*) = tensor, *Q*^MSD^(*t*): ⟨Δ*r*_*i*_(*t*)Δ*r*_*j*_(*t*) ⟩, and constructed a normalized anisotropy, 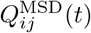:

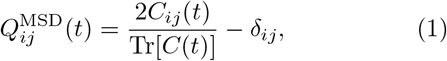

where *δ*_*ij*_ is the two-dimensional Kronecker delta. From this tensor, we extracted a scalar order parameter, *λ*(*t*), defined as its positive eigenvalue. This parameter directly measures the directional asymmetry of diffusion, bounded between *λ* = 0 for perfectly random, isotropic motion and *λ* = 1 for strictly one-dimensional transport.

As shown in Fig. 3B, pre-stressed mucus exhibits a significantly higher order parameter (*λ*) than unstressed controls, confirming that our macroscopic observations reflect a genuine structural realignment at the microscopic scale. The time dependence of *λ*(*t*) reveals how particles navigate this altered landscape over different time lags. At short times, the tracer particles probe local, unconstrained fluid voids and exhibit near-isotropic motion (*λ*≈0). At longer times, however, as the particles diffuse far enough to encounter the boundaries of the aligned mucin mesh, their trajectories become increasingly channeled along the axis of applied stress, causing *λ*(*t*) to rise. The steady-state value of *λ* thus captures the orientational symmetry of the sheared network, governed by both the degree of fiber alignment and the size of the particle relative to the mesh spacing.

By analyzing the principal eigenvectors of 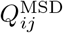, we determined the exact direction of this maximum transport bias (Fig. 3C). The alignment direction is slightly offset from the primary shear axis, a characteristic of polymer networks reorganizing under high-Weissenberg number conditions [24, 25] that we estimate for ciliary forcing in Section C. Together, these microscopic findings demonstrate that the mechanical memory of the mucin network creates preferential structural pathways. By biasing particle mobility along the axis of past deformation, this structural memory provides a physical mechanism that could actively guide the transport of inhaled pathogens and therapeutic vehicles within the lungs.

### C. A separation of bond relaxation timescales underlies the mechanical memory

The persistence of the mechanical asymmetry established in our experiments raises a fundamental question: what structural feature allows the mucin network to retain a directional memory of applied stress? The answer lies in the relaxation behavior of the network across widely separated timescales. Stress relaxation experiments (Fig. 2D) reveal a multi-stage decay profile. For instance, following an initial deformation, the network stress drops rapidly from approximately 38 Pa to 7 Pa within the first 30 s, before entering a decay phase that spans several hours. This two-stage relaxation cannot be explained by a single-timescale viscoelastic model. Instead, it points to a dual-subnetwork architecture comprising a fast network of transient associations—such as ionic bridges, hydrogen bonds, and hydrophobic interactions [13]—that rapidly break and reform under load, alongside a slow scaffold of long-lived physical entanglements and covalent disulfide bonds that reorganizes only over much longer timescales [26]. While the fast network dictates the short-term viscoelastic response and immediate structural plasticity, the slow scaffold maintains overall network integrity and, crucially, preserves a directional imprint of the deformation long after the transient bonds have re-equilibrated.

While native mucus exhibits a continuous hierarchy of relaxation timescales rather than two discrete regimes [16], our dual-subnetwork formulation serves as a minimal, coarse-grained representation of this spectrum. Despite its simplicity, this model captures the network’s structural memory.

To provide a mechanistic and quantitative basis for this picture, we employ the transient network theory (TNT) [18, 27], in which the mucus is modeled as two parallel subnetworks whose bonds attach and detach with kinetic rates *k*_*a*_ and *k*_*d*_, respectively. In this framework, the total deformation gradient ***F*** is decomposed multiplicatively into an elastic part ***F***_*e*_ and an inelastic part ***F***_*p*_, as ***F*** = ***F***_*e*_ · ***F***_*p*_, where ***F***_*p*_ accounts for the stress-free reference configuration of newly attached bonds and ***F***_*e*_ captures the elastic stretch and rotation of the network fibers relative to that reference. The elastic left Cauchy– Green deformation tensor is then defined as 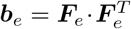, and its trace Tr(***b***_*e*_) measures the elastic stretch of the network, with Tr(***b***_*e*_) = 3 corresponding to the stress-free state. Both subnetworks are assumed to deform under the same total deformation gradient ***F***, analogous to the standard linear solid model, but each has its own elastic component 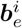 governed by its own bond kinetics. The time evolution of 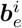 for each subnetwork is [17]:

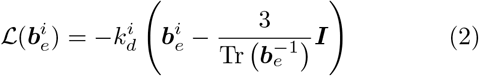

where 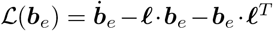 is the Oldroyd rate (also referred to as the Lie time derivative) of the spatial tensor 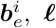 is the velocity gradient tensor, ***I*** is the identity tensor, and the superscript *i* = *s* or *f* denotes the slow and fast subnetworks respectively. The right-hand side drives 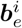 toward its stress-free isotropic state at a rate set by 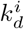: when bonds detach and reattach, the newly formed bonds adopt the current configuration as their reference, progressively releasing the stored elastic strain. The total strain energy density per unit reference volume is 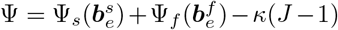, where *κ* is a Lagrange multiplier enforcing incompressibility. Given the strain-stiffening observed in our experiments, we adopt the Gent strain energy density function for both subnetworks to capture material and geometrical nonlinearities:

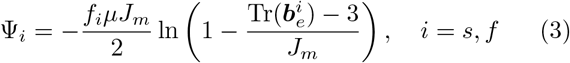

where *f*_*i*_ denotes the network fraction, with *f*_*s*_ = 1−*f* for the slow subnetwork and *f*_*f*_ = *f* for the fast subnetwork, *µ* is the total shear modulus, and *J*_*m*_ is the Gent non-linearity parameter.The resulting Cauchy stress tensor is 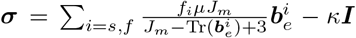. The detachment rate *k*_*d*_ is not constant but increases with the elastic stretch of the fibers, consistent with our stress relaxation data (Fig. 2D). Following recent studies of stress-dependent bond kinetics [28, 29], we model the detachment rate of the fast network using the Ellis form [30]:

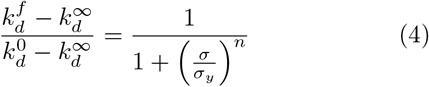

where 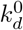 and 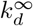 are the limiting detachment rates at low and high stress, respectively, *σ*_*y*_ is a critical stress threshold, and *n* controls the width of the transition (Fig. 4A). The slow network detachment rate 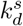 is taken as stress-independent and small, consistent with the slow relaxation timescale observed experimentally.

**FIG. 4.**
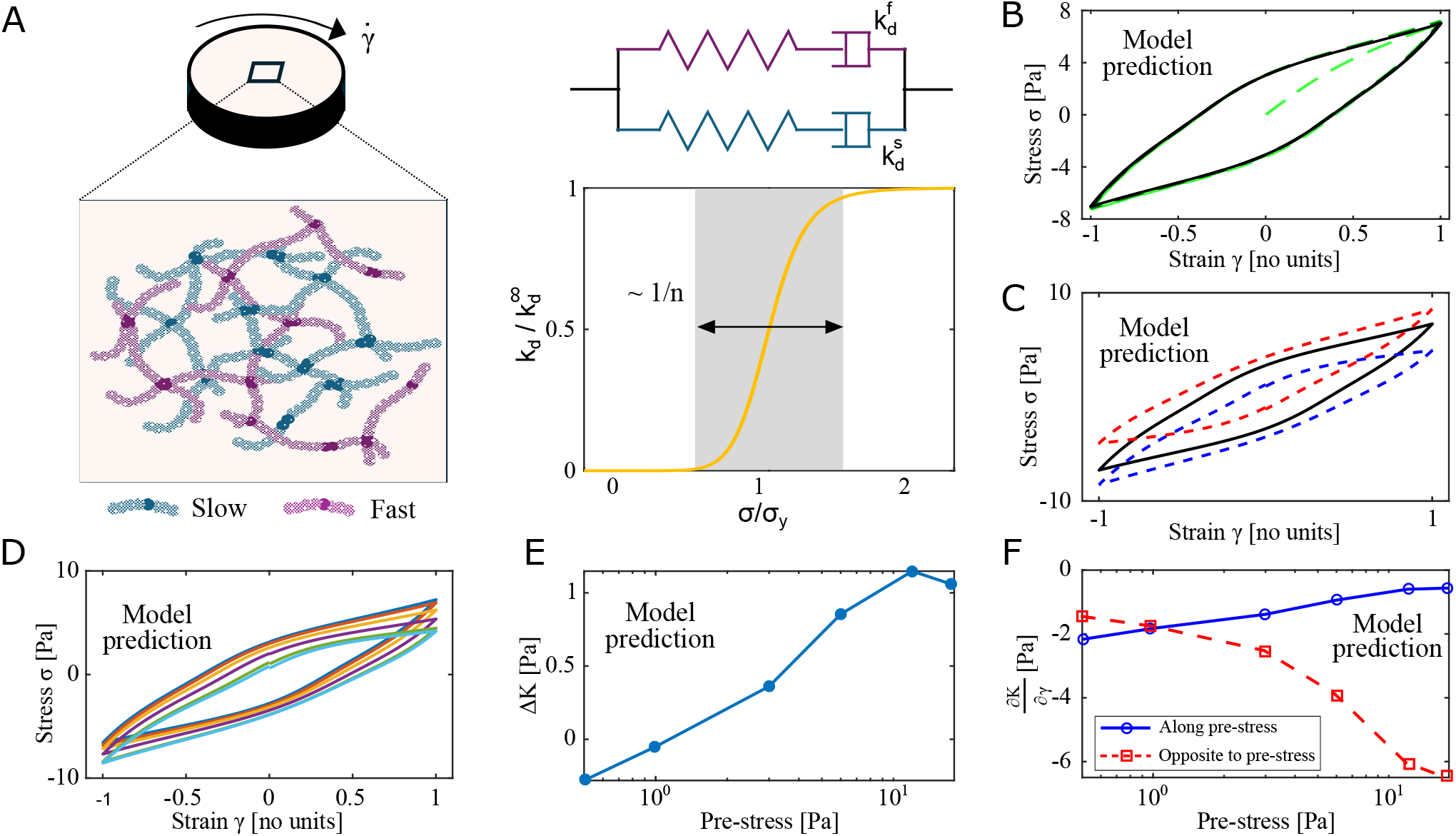
Model and predictions. (A) A schematic illustrating the underlying mucus network during shear rheology testing. The network consists of two crosslinked subnetworks: one characterized by a fast bond detachment rate, 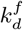, and another composed of bonds with significantly longer lifetimes, characterized by 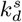. The diagram illustrates the dependence of the bond detachment rate, 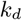, on the applied stress. (B) - (C) are the qualitative model predictions for the experimental results of Fig. 1, and (D) to (F) reproduce the results of Fig. 2 qualitatively.

Given the substantial sample-to-sample variability in mucus rheology, we do not seek an exact quantitative fit but rather demonstrate that the model reproduces the experimentally observed trends within the range of measured values. The model captures the direction-dependent shift of the hysteresis loop after pre-stressing (Fig. 4B–D), the monotonic growth of Δ*K* with prestress magnitude (Fig. 4E), and the transition from strain stiffening to strain softening in the direction opposite to the pre-stress (Fig. 4F), using parameters *k*^0^ = 0.002 s^*−*1^, 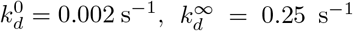, *σ*_*y*_ = 3 Pa, *n* = 2, 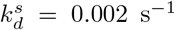, *f* = 0.6, *J*_*m*_ = 12.0, and *µ* = 10 Pa. The model provides a clear mechanistic interpretation of the observed memory. Under pre-stress, the stress-dependent detachment rate 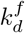 accelerates the rupture of transient associations in the fast network, facilitating local chain alignment. As these bonds reform in the deformed configuration, the slow scaffold, whose relaxation time far exceeds the duration of the experiment, preserves a non-equilibrium, anisotropic state even after the macroscopic stress has vanished. The network energy minimum is therefore reached in a configuration that retains the directional imprint of the applied load, producing the residual mechanical asymmetry observed in Section A.

Mechanical hysteresis and direction-dependent nonlinear elasticity arising from pre-stress have been observed previously in crosslinked biopolymer networks, most directly in reconstituted networks of semiflexible actin filaments, where shear training induces persistent asymmetry in the nonlinear response [20, 31]. Strain-stiffening and shear-induced anisotropy, including negative normal stresses under shear, have likewise been documented in other semiflexible biopolymer networks such as fibrin and collagen [32, 33]. The mechanisms invoked in those systems typically involve the mechanical properties of the constituent filaments and their alignment under stress. The mucin network is composed of flexible, heavily glycosylated chains rather than semiflexible filaments, and the mechanism we identify here is correspondingly distinct: the persistence of the directional response after the load is removed arises at the network level, from the separation of timescales between fast transient bonds and a slow long-lived scaffold. This composite-network mechanism may apply more broadly to viscoelastic materials whose architecture provides a separation of bond lifetimes, independent of the rigidity of individual constituents.

This picture has direct implications for the physiological regime of ciliary transport. The slow scaffold relaxation time, on the order of hours as shown by the stress relaxation data, far exceeds the period of ciliary beating (*T* ≈ 0.05 s to 0.2 s at 5 Hz to 15 Hz). The Deborah number *De* = *λ/T*, comparing the slow relaxation time *λ* to the ciliary period, therefore lies in the range *De* ≈ 10^4^–10^6^, well beyond the regime where the network can relax between strokes. Similarly, the Weissenberg number 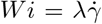, with shear rates 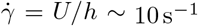 to 60 s^*−*1^ estimated from typical mucus transport velocities *U* ≈ 100 µm s^*−*1^ to 300 µm s^*−*1^ and layer thickness *h* ≈ 5 µm to 10 µm, yields *Wi* ≈ 10^3^–10^5^, placing ciliary transport deep in the nonlinear elastic regime. Together, these dimensionless numbers indicate that successive ciliary strokes do not allow sufficient time for the slow scaffold to relax, so directional deformations compound across cycles rather than canceling, progressively building the structural memory and anisotropy quantified in Sections A and B.

### D. Physiological relevance

The experiments presented above were performed under controlled laboratory conditions using sustained and cyclic pre-stresses of defined magnitude and duration. A central question is whether the stress magnitudes required to produce measurable mechanical memory are representative of those mucus experiences in the living lung. We address this by estimating the directional mechanical loads imposed on the mucus layer by different physiological processes and mapping them onto the anisotropy threshold established in Section A. The resulting comparison (Fig. 5) places ciliary beating just below the yield stress threshold established by our experiments, while coughing exceeds it, suggesting that the two principal clearance mechanisms operate in mechanically distinct regimes.

**FIG. 5.**
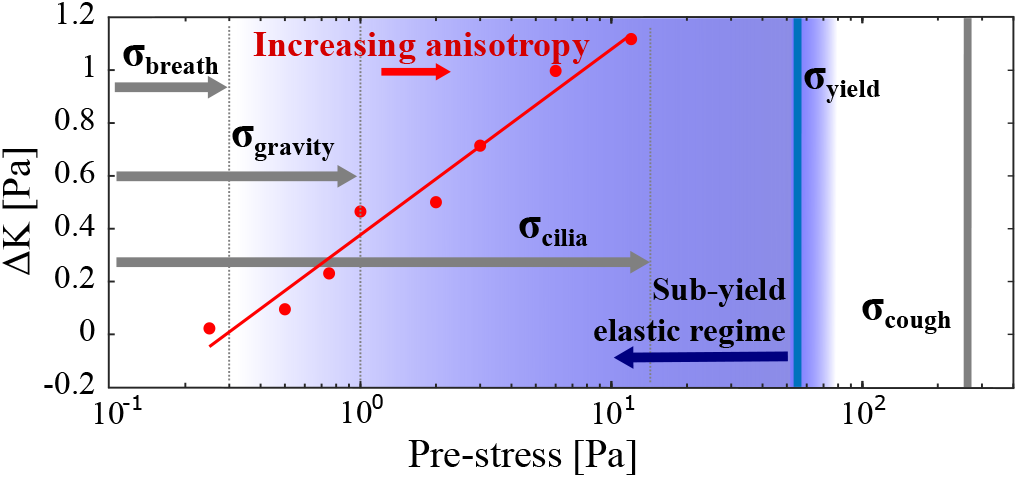
Mapping of physiological airway stresses onto the mechanical anisotropy curve (reproduced from Fig. 2B). While tidal breathing falls below the threshold for structural alignment, gravitational loading and ciliary metachronal activity generate stresses sufficient to drive persistent anisotropy (Δ*K*). Extreme events like coughing drive mucus far past the yield threshold into bulk clearance.

Mucus in the respiratory tract experiences a wide range of directional mechanical loads depending on the physiological process involved. During tidal breathing, air-flow generates a nearly constant shear stress of approximately 0.045 Pa across most airway diameters [4, 6], although stresses in the nasal cavity can reach 0.3 Pa [34]. These values fall below the threshold at which measurable anisotropy develops in our experiments (Fig. 5), suggesting that breathing alone is insufficient to persistently reorganize the mucin network. Gravitational loading of a vertical mucus layer of thickness *H* produces a shear stress *σ*_gravity_ = *ρgH* at the mucus–cilia interface. For *H* ∼ 10 µm to 100 µm and *ρ* ∼ 10^3^ kg m^*−*3^, we estimate *σ*_gravity_ ∼ 0.1 Pa to 1 Pa, corresponding to a modest but nonzero anisotropy Δ*K* ∼ 0.4 (Fig. 3D). This persistent gravitational stress could therefore prime the network for further alignment under additional loading.

The most significant source of directional stress is the coordinated beating of cilia in metachronal waves. By modeling the kinematics of cilium tip trajectories (Supplementary Information), we estimate the local strain imposed on the mucus between cilia separated by half a metachronal wavelength to oscillate between 0.5 and 1.5 for representative human airway parameters [35]. Given a Young’s modulus *E* ∼ 10 Pa, this generates a local stress *σ*_cilia_ ≈15 Pa, which corresponds to a high degree of anisotropy (Δ*K* ∼ 1; Fig. 5). Coughing represents the most extreme case, with theoretical models suggesting wall shear stresses as high as 170 Pa [5], far into the strongly anisotropic regime and the yield stress (Fig. 5). These estimates collectively suggest that the lung operates across a wide range of the anisotropy curve established in our experiments, with ciliary activity alone sufficient to transform mucus into a directionally aligned medium.

The emergence of shear-induced mechanical memory may have important consequences for the efficiency of mucociliary transport. In an isotropic network, the energy expended by a cilium during its power stroke is distributed uniformly through the gel. If the mucin network progressively aligns along the axis of the power stroke (as our results suggest it would under sustained ciliary stress), the resulting directional stiffening could facilitate long-range mechanical coupling between neighboring cilia along the direction of mucus alignment, while cilia would be less mechanically coupled to their neighbors orthogonal to the mucin alignment direction. The feedback between ciliary forcing and network structure would imply that mucociliary transport is not merely a problem of fluid propulsion but involves a dynamic co-organization of the material being transported. In pathological states where the network is overly dense or crosslinked (e.g., cystic fibrosis or chronic obstructive pulmonary disease (COPD)), this alignment could become too rigid and contribute to excessive mechanical loading that hinders mucus transport.

The directional bias in particle diffusion established in Section C adds a further layer of physiological consequence. The preferential diffusion pathways created by network alignment could guide the transport of inhaled pathogens along the axis of ciliary stress, consistent with prior observations that stress-aligned mucin networks rectify bacterial trajectories from random walks into parallel paths [7]. The size dependence of the cage anisotropy suggests that this effect may sort pathogens by size, with particles comparable to the mesh spacing experiencing stronger directional bias than larger ones. This raises the possibility that the structural memory of the mucus network plays an active role in pathogen capture and directional clearance, rather than acting merely as a passive sticky trap. Finally, we note that the stress magnitudes generated by ciliary beating are of the same order as the yield stress of mucus (*σ*_*y*_ ∼ 55 Pa, Fig. 5), which in our experiments separates the elastic regime from irreversible flow. This proximity to the yield threshold may not be coincidental. A material operating just below its yield stress can accumulate structural alignment under repeated directional loading while maintaining the coherent layer geometry required for ciliary transport. Stresses above yield, such as those generated during coughing, would instead drive irreversible flow and bulk clearance. This suggests that the mechanical architecture of mucus, its yield stress, its dual relaxation timescales, and its capacity for directional memory, may be collectively tuned to support two distinct clearance mechanisms: continuous ciliary transport in the sub-yield elastic regime, and episodic cough clearance above yield. We emphasize that this interpretation is based on order-of-magnitude stress estimates and requires direct in vivo validation, but it provides a physically grounded framework for future investigations of mucociliary function in health and disease.

## CONCLUSION

We showed that lung mucus develops a persistent directional mechanical anisotropy when subjected to stresses of physiologically relevant magnitude, and that this anisotropy reflects a geometric reorganization of the mucin network that biases microscopic particle transport. These findings rest on three interlocking results: bulk rheology establishes that directional pre-stress imprints a mechanical asymmetry that persists after the load is removed; a transient network model with dual relaxation timescales explains this persistence as a structural memory encoded in the slow scaffold of the network; and particle-tracking microrheology confirms that this memory reorganizes the network geometry at the scale of biological particles, creating preferential diffusion pathways along the axis of applied stress.

The central conceptual contribution of this work is to reframe mucus not as a passive isotropic barrier, but as a material with directional mechanical memory. Its response to stress is not determined solely by its instantaneous state but by its history of loading, which is written into the slow scaffold of the mucin network and expressed as a directional bias in both mechanical stiffness and particle mobility. The transient network model developed here provides a quantitative bridge between this macroscopic behavior and the underlying molecular bond kinetics, identifying the stress-dependent detachment rate of transient associations as the key parameter controlling how rapidly memory accumulates and how long it persists.

The physiological implications of these results are significant, but require further investigation. Our stress estimates suggest that ciliary beating, gravitational loading, and coughing place the lung across a wide range of the anisotropy curve we have characterized, with ciliary activity alone sufficient to drive substantial network alignment. The proximity of ciliary stresses to the mucus yield threshold raises the possibility that the mechanical architecture and dynamics of mucus is collectively tuned to support two distinct clearance mechanisms: continuous directional transport in the sub-yield elastic regime, and episodic bulk clearance during coughing. Whether this mechanical memory is a hallmark of healthy mucociliary function or becomes disrupted in obstructive airway diseases such as cystic fibrosis and chronic obstructive pulmonary disease remains an open and important question. More broadly, the framework developed here provides a physically grounded basis for future in vivo studies of mucus mechanics, pathogen navigation, and the rational design of inhaled therapeutic particles.

## ACKNOWLEDGEMENTS

This work was supported by the Boettcher Foundation’s Webb-Waring Biomedical Research Program (N.F.M.), the National Science Foundation Grant No. 2527013 (N.F.M.), the University of Colorado ABNexus Award (N.F.M.), the NIH/CU Molecular Biophysics Program (A.G.P. and A.G.G.), and the NIH Biophysics Training Grant T32 GM145437 (A.G.G.).

## METHODS

### 1. Extracting cow tracheal mucus

Cow lungs and trachea were obtained from Elizabeth Meat Lockers in Elizabeth, CO. The trachea was carefully scraped using a spatula to get the mucosal material out within an hour of butchering. Up to 5 ml of mucus was obtained and collected in a centrifuge tube. Afterwards, the trachea was removed and the lungs were cut along the bronchi using scissors. A smaller amount (under 1 ml) was extracted from the bronchi. The mucus was stored at 4^*°*^C and used for up to 3 days.

### 2. Rheology on cow tracheal mucus

An Anton Paar MCR302e Rheometer was used to perform rheology experiments on mucus in the parallel plate configuration. A 50 mm diameter plate was used. About 300 *µ*l of mucus was used per experiment, and the plate distance was set to 100 *µ*m. The sample was enclosed in a chamber that maintained a temperature of 25^*°*^, and a humidity level of 95% to prevent water evaporation. To obtain stress *σ*–strain *γ* curves at different pre-stress levels, each pre-stress was applied and held for 250 s, after which the corresponding curve was measured. A 5 min recovery period separated successive measurements to allow relaxation before applying the next pre-stress.

