## Supplemental Information for "Stress-Induced Mechanical Memory in Respiratory Mucus: Anisotropy, Network Reorganization, and Directional Transport"

(Dated: July 16, 2026)

### VISCOELASTIC MEASUREMENTS ON COW LUNG MUCUS

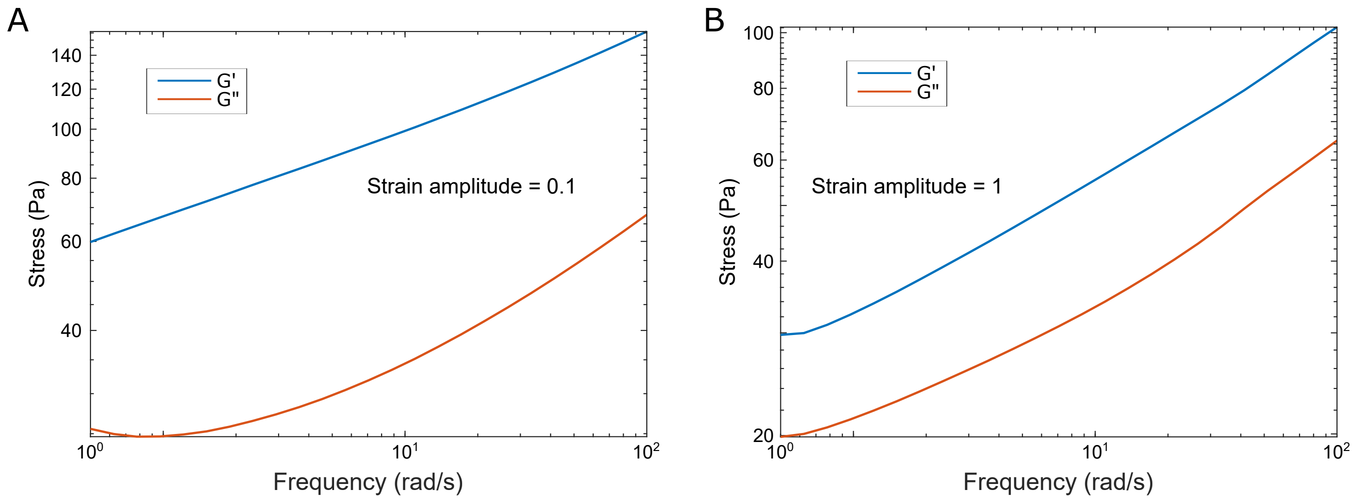


FIG. S1. Viscoelastic measurements on cow lung mucus at two strain amplitudes. (A) Storage modulus $G'$ and loss modulus $G''$ as functions of oscillation frequency for a strain amplitude of 0.1. (B) Storage modulus $G'$ and loss modulus $G''$ as functions of oscillation frequency for a strain amplitude of 1. In both measurements, the storage modulus dominates the loss modulus.

### DETERMINING THE YIELD STRAIN RATE IN COW LUNG MUCUS

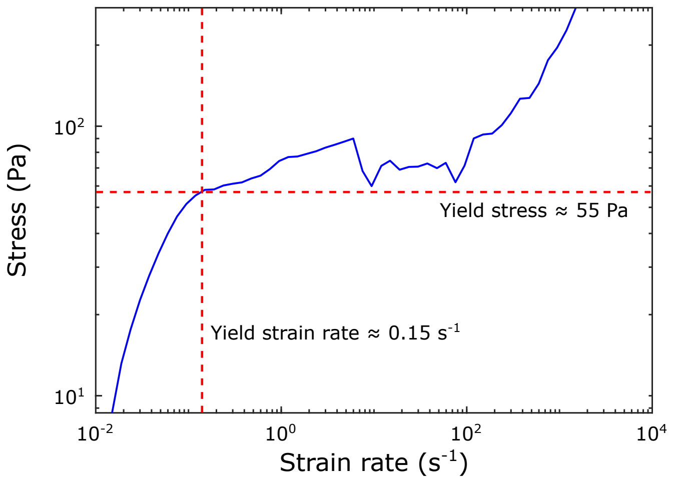


FIG. S2. Determining the yield strain rate i.e. the maximum strain rate for elastic deformation in a cow lung mucus sample. As seen in the plot, the stress increases linearly with strain rate at low values. However, at approximately 0.15 Hz (defined as the yield strain rate), the stress begins to plateau and exhibit erratic fluctuations. The stress at the yield strain rate (55 Pa) was defined as the yield stress of the mucus sample. This behavior indicates that the sample is undergoing irreversible deformation beyond this point. Hence, all hysteresis measurements were performed at strain rates lower than the yield strain rate.

### STRAIN RATE ON COW LUNG MUCUS DURING PRE-STRESS

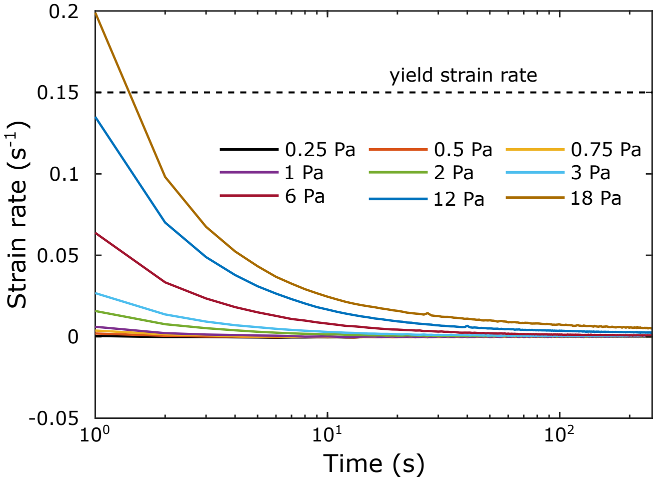


FIG. S3. Strain rates on cow mucus during pre-stress. Apart from a few seconds in the beginning for pre-stress >12 Pa, the strain rates are always lower than 0.15 $s^{-1}$, which was determined earlier to be the Yield strain rate.

### HYSTERESIS MEASUREMENTS ON SEVERAL COW LUNG MUCUS SAMPLES

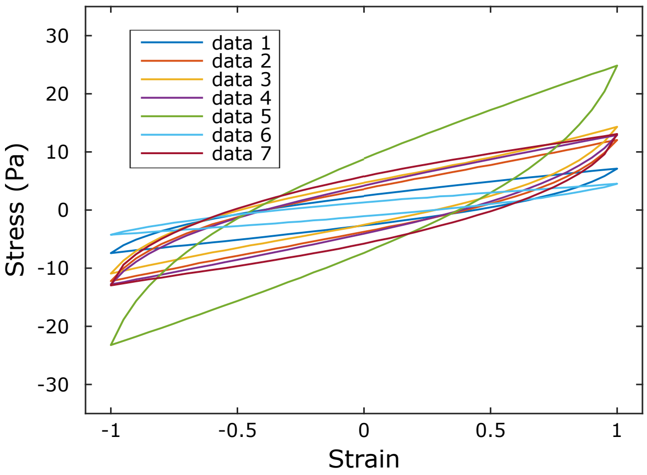


FIG. S4. Variability of the stress–strain response in cow mucus. Stress–strain measurements for seven cow mucus samples show modest amplitude variations, likely reflecting differences in water content. The consistent qualitative form of the stress–strain loops across samples indicates an identical underlying network structure.

### ANALYSIS OF DIRECTIONAL DIFFERENTIAL MODULUS

To characterize the emergence of mechanical anisotropy, we analyzed the differential shear modulus, $K(\gamma) = \mathrm{d}\sigma/\mathrm{d}\gamma$, as a function of both the instantaneous strain and the magnitude of the applied pre-stress.

Along the pre-stress direction ($\gamma < 0$), the mucin network exhibits a constant resistance to deformation. As shown in Fig. S5A, $K(\gamma)$ initially fluctuates for small strains before reaching a nearly constant value. Notably, at the maximum strain along this direction ($\gamma = -1$), the differential modulus becomes largely independent of the pre-stress magnitude. This suggests that the pre-stressing phase leads to maximum possible alignment of the mucin fibers. Any further loading in the same direction encounters a consistently reinforced structural scaffold.

In contrast, the response in the direction opposite to the pre-stress ($\gamma > 0$) indicates a progressive softening of the network. As shown in Fig. S5D, the differential modulus remains nearly constant across the strain range within a single cycle but decreases monotonically with increasing pre-stress magnitude. This behavior is consistent with a structural "loosening" of the network when the load is reversed. The fibers aligned during pre-stress likely undergo buckling or rapid detachment of transient crosslinks, significantly reducing the network's ability to resist opposing shear.

The divergence between Fig. S5A and Fig. S5B suggests a molecular-scale structural "polarization" of the mucin network. In the pre-stress direction, fibers are pulled into tension, where they resist further deformation through high-modulus entropic or enthalpic stretching. In the reverse direction, these now-aligned fibers are subjected to compression, potentially leading to fiber buckling or the forced dissociation of weak, transient crosslinks. This directional asymmetry creates a material that remains compliant in one direction while providing reinforced resistance against the other.

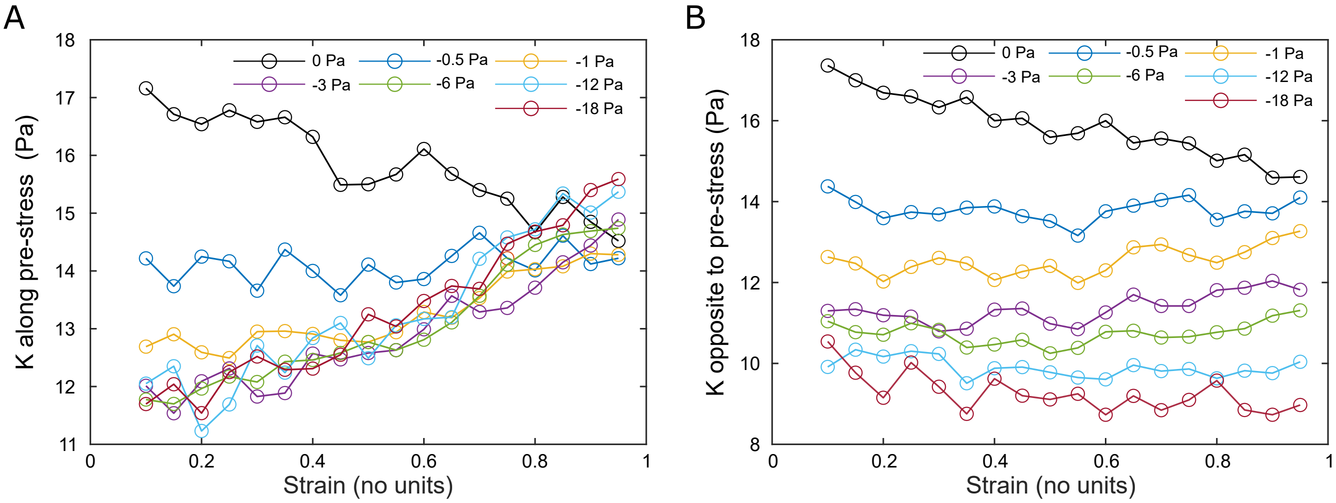


FIG. S5. Directional differential modulus vs strain rate for several magnitudes of pre-stress. (A) Differential stress modulus in the direction of the applied pre-stress from 0 to -18 Pa. (B) Differential stress modulus opposite to the direction of the applied pre-stress from 0 to -18 Pa.

### RHEOMICROSCOPY SETUP

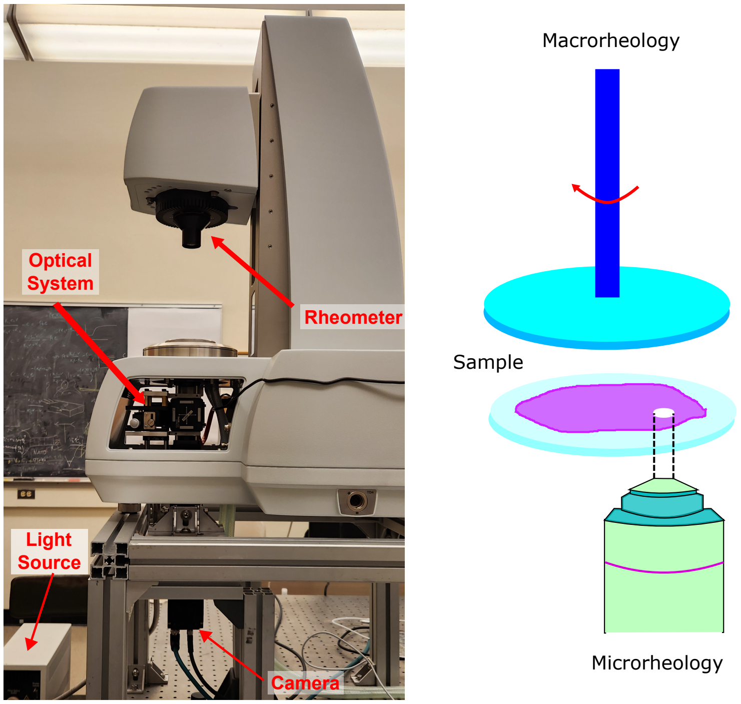


FIG. S6. Rheomicroscopy setup. Below the bottom plate of the rheometer, we integrated an optical imaging system that allows direct visualization of the sample during microrheology measurements. This enables us to observe particles diffusing within the sample in real time while the sample is being sheared. In turn, the setup provides a direct view of the microrheological response of the material to applied shear stress by tracking the motion of embedded particles.

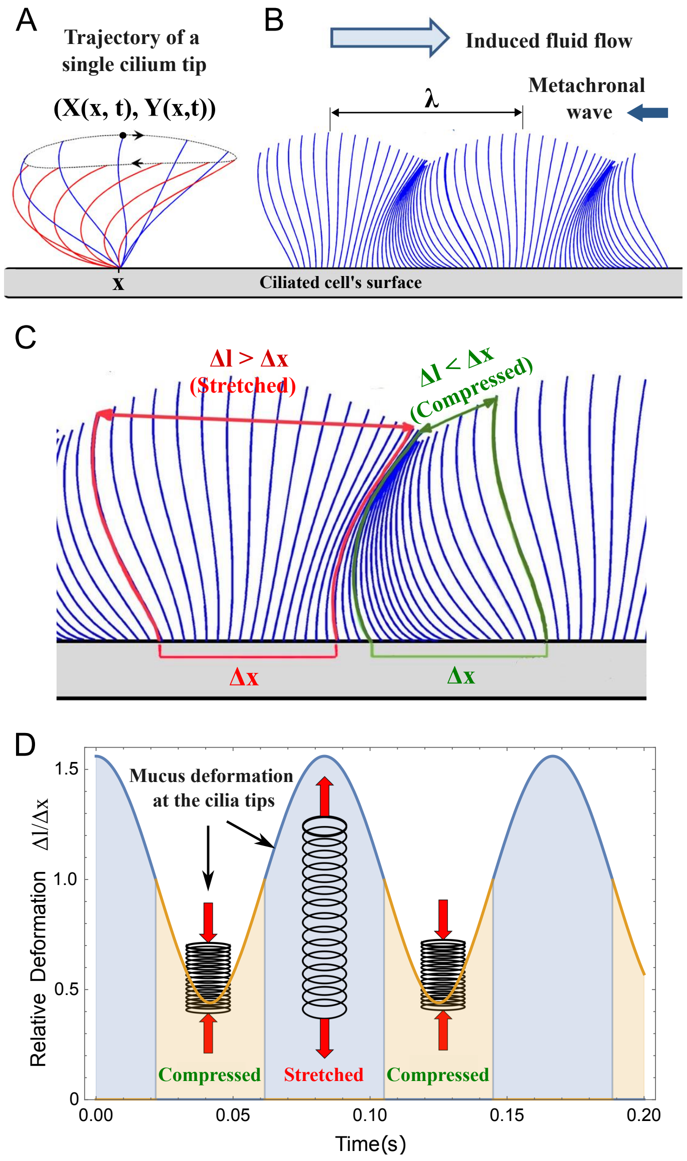


FIG. S7. Sketch of a metachronal wave in a ciliary tissue and associated mucus deformation. (A) Trajectory of a single cilium tip moving periodically. (B) Snapshot of the ciliary configuration at a given time. Image adapted from [1]. (C) Distance between cilia tips $\Delta l$ corresponding to similar distances of the cilia base $\Delta x$. Cilia tip distances depend on base position for a given time. (D) Relative deformation of mucus at the cilia tips for cilia separated a distance of half the metachronal wavelength. The cilia tips are periodically pulled apart and brought close, inducing periodic stretching and compression of mucus.

### KINEMATIC MODEL FOR CILIA STRETCHING OF MUCUS

The coordinated beating of cilia in metachronal waves imposes cyclic deformations at the mucus-cilia interface. We model the trajectory of a single cilium tip as an ellipsoid of coordinates $(X, Y)$:

$$X(x,t) = x - a\cos\left[\frac{2\pi x}{\lambda} + \frac{2\pi t}{T}\right], \tag{1}$$

$$Y(x,t) = b\,a\sin\left[\frac{2\pi x}{\lambda} + \frac{2\pi t}{T}\right], \tag{2}$$

where $x$ is the coordinate of the cilium base, $a$ and $b$ are the semi-axes of the ellipse, and $\lambda$ and $T$ are the wavelength and period of the metachronal wave, (Fig. S7A) [1]. From these kinematics, we estimate the separation $\Delta l$ between the tips of two cilia whose bases are separated by a distance $\Delta x$ (Fig. S7C). This network deformation reaches its maximum intensity when the distance between the bases corresponds to half the metachronal wavelength ($\Delta x = \lambda/2$), as shown in Fig. S7D. Using representative values for human airways—$a = 7\,\mu\text{m}$, $b = 1$, $T = 0.083\,\text{s}$ (12 Hz), $\lambda = 50\,\mu\text{m}$, and $\Delta x = 25\,\mu\text{m}$ [2]—the relative compression and extension ($\Delta l/\Delta x$) oscillates between 0.5 and 1.5. With a maximum strain estimate of $\gamma \approx 1.5$ and a Young's modulus $E \sim 10\,\text{Pa}$, we estimate a stress $\sigma_c = E\gamma \sim 15\,\text{Pa}$. This stress yields $\Delta K \sim 1$ (Fig. 2B), indicating that ciliary activity alone is sufficient to transform mucus into an anisotropic medium.

[1] M. Bottier, M. Peña Fernández, G. Pelle, D. Isabey, B. Louis, J. B. Grotberg, and M. Filoche, PLoS computational biology **13**, e1005552 (2017).
[2] A. Burn, M. Schneiter, M. Ryser, P. Gehr, J. Rička, and M. Frenz, European Biophysics Journal **51**, 51 (2022).
